# A semi-parametric multiple imputation method for high-sparse, high-dimensional, compositional data

**DOI:** 10.1101/2024.09.05.611521

**Authors:** Michael B. Sohn, Kristin Scheible, Steven R. Gill

## Abstract

High sparsity (i.e., excessive zeros) in microbiome data, which are high-dimensional and compositional, is unavoidable and can significantly alter analysis results. However, efforts to address this high sparsity have been very limited because, in part, it is impossible to justify the validity of any such methods, as zeros in microbiome data arise from multiple sources (e.g., true absence, stochastic nature of sampling). The most common approach is to treat all zeros as structural zeros (i.e., true absence) or rounded zeros (i.e., undetected due to detection limit). However, this approach can underestimate the mean abundance while overestimating its variance because many zeros can arise from the stochastic nature of sampling and/or functional redundancy (i.e., different microbes can perform the same functions), thus losing power. In this manuscript, we argue that treating all zeros as missing values would not significantly alter analysis results if the proportion of structural zeros is similar for all taxa, and we propose a semi-parametric multiple imputation method for high-sparse, high-dimensional, compositional data. We demonstrate the merits of the proposed method and its beneficial effects on downstream analyses in extensive simulation studies. We reanalyzed a type II diabetes (T2D) dataset to determine differentially abundant species between T2D patients and non-diabetic controls.

## 1 Introduction

Over the past decade, the US National Institutes of Health has invested more than one billion dollars in human microbiome research, and this investment has confirmed the influence of the microbiome on human health and disease (Proctor, 2019). The dysbiosis of the microbiome has been shown to contribute to the pathogenesis of a range of diseases, including oral diseases, metabolic disorders, autoimmune diseases, and neurodevelopmental disorders (Wade, 2013; Vetizou *et al*., 2015; Warner, 2019). This wealth of discoveries from human microbiome research has been made possible due to high-throughput sequencing technologies (Li, 2015), which allow us to study microbes by analyzing all genomic contents directly obtained from environmental samples without prior cloning and culturing individual microbes.

The result of preprocessing the sequenced microbiome samples is a table of counts (called a taxonomic profile), where rows and columns typically represent samples and microbes (or taxa), respectively. Figure 1 shows a typical taxonomic profile of microbiome data. The left box plot represents a variation in the total number of taxa (i.e., sequencing depth) in microbiome samples. The right heatmap represents the abundance of taxa, where the white color indicates the absence of taxa, and the gray scale indicates a level of the abundance of taxa in the samples: darker gray, higher abundance. Because of large variations in sequencing depths, as shown in Figure 1, observed counts of taxa are typically transformed into proportions, making the taxonomic profile into compositional data. Moreover, as indicated by a high portion of white color in the heatmap, microbiome data contain excessive zeros.

**Figure 1:**
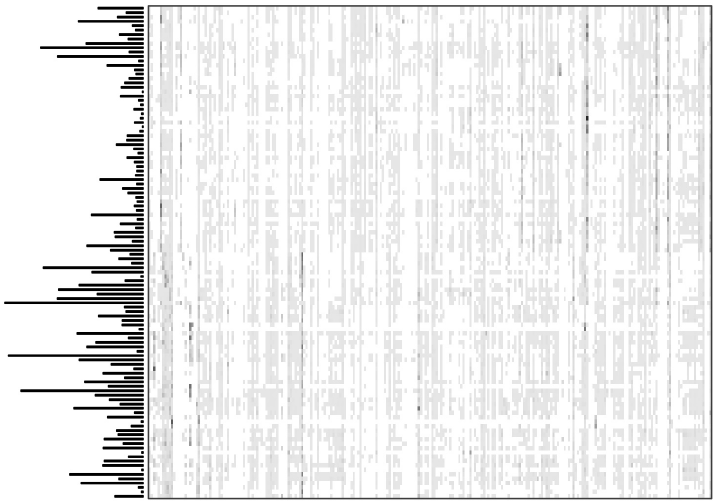
Typical characteristics of microbiome data. The left plot represents a variation in the total numbers of microbial features. The right heatmap represents the abundance of microbial features.(e.g., 0.5).

As zeros are undefined in log-ratio, which is the foundation of compositional data analysis (Aitchison, 1986), zeros must be replaced with some positive values. The simplest but most common approach is to replace zeros with a small value (e.g., 0.5). However, the choice of the small value may have substantial effects on downstream analysis and cannot be theoretically or empirically justified. Some remedial approaches are model-based multiplicative lognormal imputation and non-parametric multiplicative simple imputation methods (Martín-Fernández *et al*., 2003). More complex imputation methods, such as simulation-based data augmentation method and K-nearest neighbor method for imputation, are also available, but they often have computational issues for highly sparse (i.e., excessive zeros) compositional data (Hron *et al*., 2010; Templ *et al*., 2016). All these imputation methods assume that zeros arise from the true absence of taxa (i.e., structural zeros) or a detection limit (i.e., technical zeros). Thus, imputed values are less than a lower limit (e.g., 0.5).

However, zeros can also originate from undetected taxa due to the stochastic nature of sampling or sampling variability (i.e., sampling zeros) (Kaul *et al*., 2017; Silverman *et al*., 2020), whose values are unconstrained to a lower limit. To address this multi-source zero problem, methods based on zero-inflated or mixture models have been proposed (Paulson *et al*., 2013; Sohn *et al*., 2015; Chai *et al*., 2018; Prost *et al*., 2021; Jiang *et al*., 2021), assuming the probability of being structural zero can be modeled with external factors. These methods may distinguish structural zeros from technical or sampling zeros to some extent if relevant genetic or external factors are included in the model. However, it is not trivial to determine which genetic or external factors may be relevant to structural zeros a priori. Even worse, the structural zero itself can arise from different sources: functional absence or functional redundancy (Manor and Borenstein, 2017), which represent opposite biological implications. One represents the absence/loss of a function, and the other represents the existence of a function performed by other taxa. Thus, distinguishing structural zeros from technical or sampling zeros may be impossible, except for a special case where zeros are in all or almost all samples of a subgroup.

This multi-source zero problem may not pose a serious problem if the proportion of structural zeros is similar across all taxa, as the mean change in proportion or ratio could be negligible. For instance, 𝔼(*m*_1_*/m*_2_) ≈ 𝔼(*m*_1_(1 − *ν*_1_)*/m*_2_(1 − *ν*_2_)) if *ν*_1_ ≈ *ν*_2_, where *m*_1_ and *m*_2_ are the mean counts of taxa 1 and 2, and *ν*_1_ and *ν*_2_ are their corresponding proportions of structural zeros. Therefore, treating all zeros as missing values or sampling zeros, except for the special case, may be less prone to substantial bias than attempting to distinguish structural zeros from technical or sampling zeros. In this manuscript, assuming this similar proportion of structural zeros, we propose a semiparametric multiple imputation method for high-sparse, high-dimensional, compositional data. We demonstrate the outperformance of the proposed method over existing methods and its beneficial effects on various downstream analyses in extensive simulation studies. We also reanalyzed a type II diabetes (T2D) dataset to determine differentially abundant species between T2D patients and non-diabetic controls.

## 2 Methods

Let *M* be an *n* × *p* matrix containing relative abundance (i.e., proportions) of taxa, *M*_−*j*_ be *M* with the *j*-th column removed, ***m***_*i*·_ be the *i*-th row vector of *M*, and ***m***_·*j*_ be the *j*-th column vector of *M*. To impute rounded zeros in the *i*-th sample, we assume that a vector of components ***m***_*i*·_, which lies in a *p* − 1 dimensional simplex space 𝕊^*p*−1^, follows a logistic normal distribution (Aitchison, 1986), which is related to a multivariate normal distribution as:

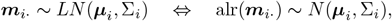

where *LN*() represents the logistic normal distribution, *N*() represents the normal distribution, *µ*_*i*_ is a *p* − 1 dimensional mean vector of log ratios, Σ_*i*_ is a (*p* − 1) × (*p* − 1) covariance matrix of log ratios for sample *i*, and alr() is the additive log-ratio transformation, which is defined as

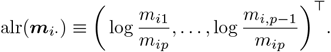

### Proposition 1

*Let* ***x*** *be a composition with p components following a logistic normal distribution with mean* ***µ*** *and covariance* Σ_*x*_, *i*.*e*., ***x*** = {*x*_1_, *x*_2_, …, *x*_*p*_} ~ *LN* (***µ***, Σ_*x*_), *and* ***y*** *be a composition with q components consisting of x*_*j*_ *and all other components in* ***x*** *amalgamated into q* − 1 *components, which also follows a logistic normal distribution, i*.*e*., ***y*** = {*x*_*j*_, *y*_2_, …, *y*_*q*_} ~ *LN* (***η***, Σ_*y*_), *where* 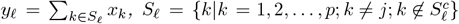, *and* 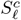 *is the complement of S*_*𝓁*_. *Then*,

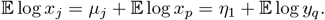

We utilize this proposition to impute sampling zeros. Specifically, for the proportions of the *j*-th taxon ***m***_·*j*_, we randomly sample *k* columns from *M*_−*j*_ without replacement and amalgamate the proportions of the randomly selected *k* columns, where *k* ∈ {1, 2, …, *p* − 1}. We repeat this random sampling with the remaining columns until the number of the remaining columns is equal to or smaller than *k*. The resultant matrix 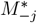 is an *n* × *q* matrix, where *q* < *p*. This approach reduces zeros in 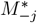 substantially when *q* ≪ *p*.

The smallest value of *q* is 2, but it completely ignores variations in other components, as we have:

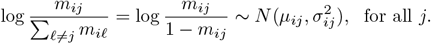

Therefore, we are only able to obtain a single imputation. To incorporate variations in other components, thus allowing for multiple imputations, we need to use *q* > 2. See Figure 2 for the graphical description of this procedure with *q* = 3, where *M* ^(*j*)^ be an *n* × 3 matrix that combines ***m***_·*j*_ with 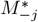.

**Figure 2:**
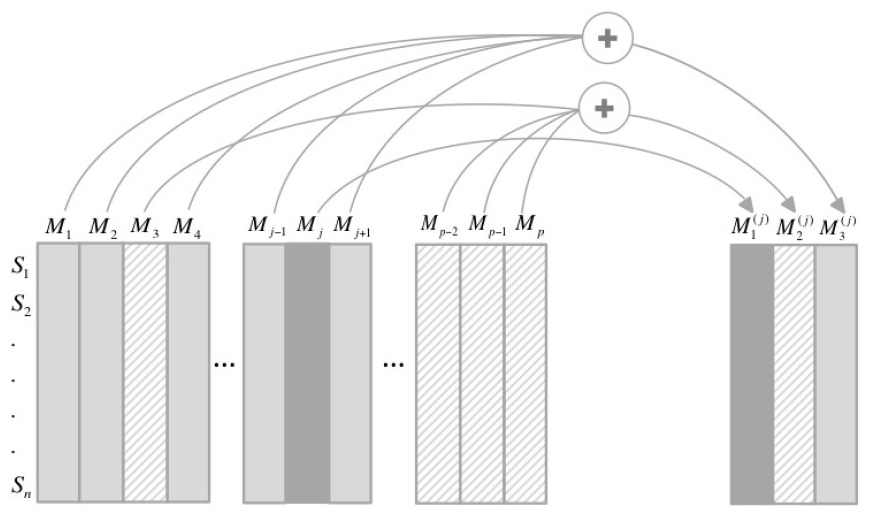
Graphical description of the proposed imputation procedure for the *j*-th column of *M*. The first colume of the resultant matrix is just the *j*-th column of *M*, the second one is the sum of ⌊ (*p*− 1)*/*2 ⌋randomly selected columns of *M*, and the third one is the sum of the remaining columns, where ⌊·⌋ is the floor function.

By the distributional assumption, i.e., 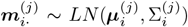, the additive log-ratio transformation of 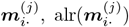 follows a multivariate normal distribution with mean 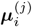 and variance 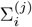. Thus, the marginal distribution of log 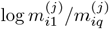 follows a normal distribution with mean 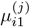 and variance 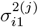.Assuming its mean can be modeled as a linear combination of covariates, i.e., 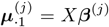, where *X* is a design matrix with the first column of 1s, and ***β***^(*j*)^ is a vector of parameters to be estimated, we can express the marginal distribution as,

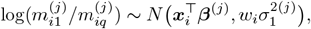

where *w*_*i*_ are sample specific values, such as rescaled sequencing depths. Therefore, we can determine the maximum weighted marginal likelihood estimator of 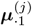 using observed values of ***m***_·*j*_ and 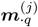, which is given by:

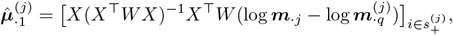

where *W* is a diagonal matrix with *w*_*i*_ on the diagonal and 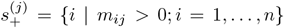. Utilizing Proposition 1, we now propose an estimator for sampling zeros:

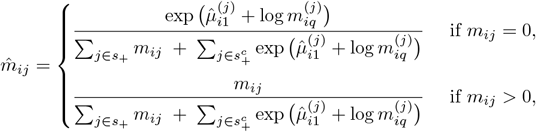

where *s*_+_ = {*j* | *m*_*ij*_ > 0; *j* = 1, …, *p*} and 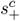 is a complement set of *s*_+_. Note that the denominator is just a rescaling factor that makes 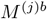_·_ sum to one. We repeat this random sampling procedure to obtain multiple *M* ^(*j*)*b*^, which are used in turn to obtain multiple 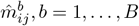. The mean of these estimates can be used to construct the final taxonomic profile table, or each estimate can be used to construct a taxonomic profile that can be used in downstream analyses. For the latter, Rubin’s rules (Rubin, 1987) or consensus rates can be used to pool estimates.

*Remarks*: For the nonzero components of composition ***m***_*i*·_, the imputed composition 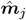 preserves the following properties:

a. Ratio invariance, i.e., 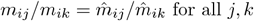
b. Subcomposition invariance, i.e., 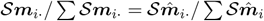, where 𝒮 is a selecting matrix, which selects a subset of the nonzero components

These invariances are a direct consequence of the multiplicative form of the proposed estimator. These also imply the invariance of algebraic operation (e.g., perturbation, power transformation) in the simplex space (Aitchison, 1986).

## 3 Results

### 3.1 Simulation Studies

We first demonstrate the performance of the proposed multiple imputation method for high dimensional, compositional data (MIC) in estimating mean proportions and ratios. We then show the effects of mismodeling structural zeros. Finally, we show its beneficial effects in some downstream analyses, specifically differential abundance analysis. The following (frequently used) methods were considered in performance comparison: replacing zeros with 0.5 (C05), multiplicative lognormal replacement (MLR) (Martín-Fernández *et al*., 2003), multiplicative simple replacement (MSR) (Martín-Fernández *et al*., 2003), and mbImpute (MBI) (Jiang *et al*., 2021). Root mean square error (RMSE) was used to measure the difference between the true and estimated values in mean proportion and ratio. The frequency of having a minimum RMSE of 200 simulations for each method was also measured.

#### 3.1.1 Accuracy

The mean proportions of compositions and mean ratios of components in compositions are primary quantities used in microbiome data analysis. Thus, we first evaluate the performance of MIC in these quantities compared to existing methods. We simulated data using zero-inflated negative binomial (ZINB) models and zero-inflated Dirichlet multinomial (ZIDM) models for this simulation study. We chose these two models to assess the robustness of the methods as they represent two opposite extreme cases: the counts of all taxa are independent of each other with ZINB, while the counts of taxa are all negatively associated with ZIDM. Model parameters were randomly selected in a setting mimicking real microbiome data, and a wide range of proportions of zeros, from 30% to 80%, was used, which covers typical proportions of zeros in real microbiome data. Details of model parameters are given in Appendix A.

Figure 3 shows the results of the proposed and existing methods in terms of mean ratios when ZINB was used with n=100 samples and p=100 taxa, and Figure 4 shows when ZIDM was used. The classical methods (i.e., C05, MLR, MSR) were heavily affected by both the correlation structure and the proportion of zeros in a composition. When the proportion of zeros was low (30%, 40%), their RMSEs were about 4-fold higher when ZIDM was used, compared to ZINB. Interestingly, as the proportion of zeros increased, their RMSEs increased when ZINB was used, whereas they decreased when ZIDM was used. Contrarily, MBI was not affected by the correlation structure. Its RMSEs were similar between ZINB and ZIDM at each proportion of zeros. However, it was greatly affected by the proportion of zeros. Its RMSEs increased as the proportion of zeros increased. It was over 4-fold higher at 80%, compared to that at 30%. On the other hand, MIC was very robust to the correlation structure. No difference in RMSE between ZINB and ZIDM was observed. It was also relatively robust to the proportion of zeros. The maximum fold difference in RMSE was less than 2. Additionally, it had the lowest mean RMSEs over 200 simulations in all settings and the highest frequency of having the lowest RMSEs in each setting. Results for mean proportions were very similar, as shown in Figures S1 and S2.

**Figure 3:**
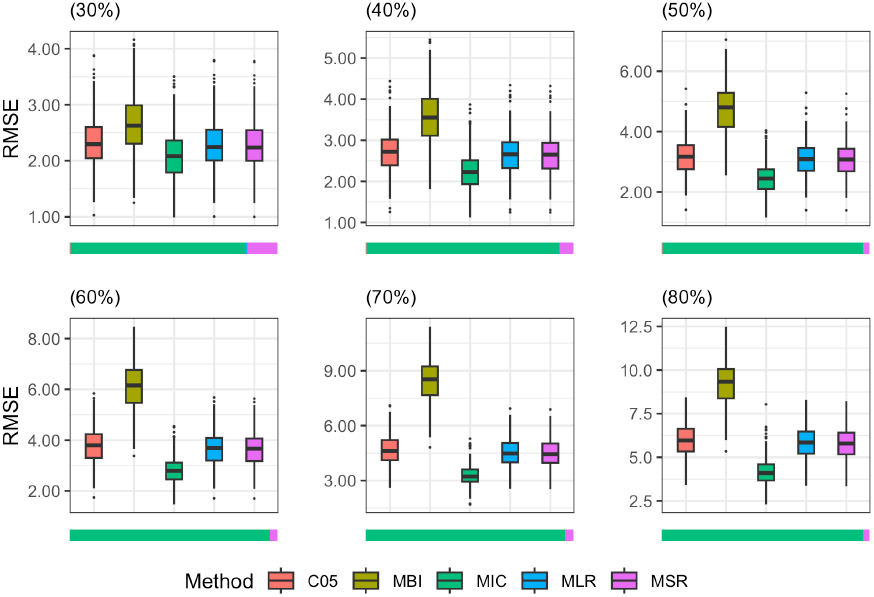
RMSE of Mean Ratios. Zero-inflated negative binomial models were used to simulate data with percent of zeros ranging from 30% to 80%. The horizontal stacked bar plots below the box plots represent the frequency of having the lowest RMSEs over 200 simulations for each method used in the comparison.

**Figure 4:**
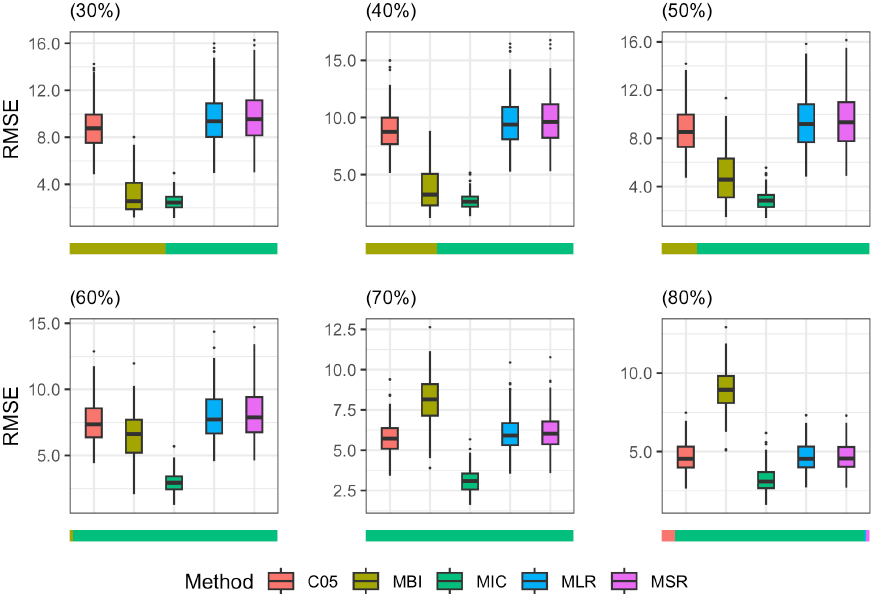
RMSE of Mean Ratios. Zero-inflated Dirichlet multinomial models were used to simulate data with percent of zeros ranging from 30% to 80%. The horizontal stacked bar plots below the box plots represent the frequency of having the lowest RMSEs over 200 simulations for each method used in the comparison.

#### 3.1.2 Effects of Mismodeling Structural Zeros on Accuracy

To investigate the effects of mismodeling structural zeros, we simulated taxonomic profiles using ZINB models with a fixed proportion of zeros for each taxon at 0.5 and a randomly generated proportion of zeros for each taxon from a uniform distribution over the interval (0.3, 0.7). MIC had a little bit higher RMSE when the proportion of zeros was randomly selected. However, the difference is not significantly different, as shown in Figure 5, supporting our premise of small effects of mismodeling structural zeros if the proportion of structural zeros is similar across all taxa. MIC with the random proportion of zeros had an even lower mean RMSE than the existing methods with the fixed proportion of zeros, as shown in Figures S3 and S4. The performance of the classical methods appreciably deteriorated when the random proportion of zeros were used, on account of constraining imputed values to a lower limit. MBI had a significantly higher RMSE in mean proportions when the proportion of zeros was randomly selected. However, it had no significant difference in mean ratios.

**Figure 5:**
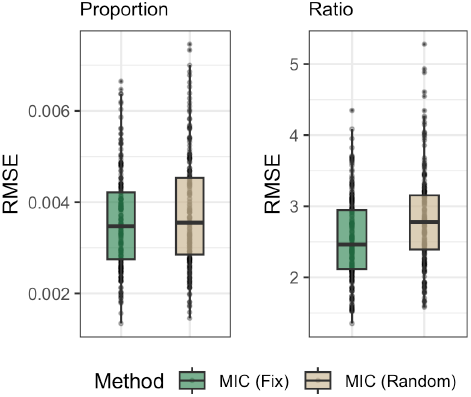
Effects of mismodeling structural zeros. Fix indicates the fixed proportion of zeros at 0.5, and Random indicates a randomly generated proportion of zeros from a uniform distribution over the interval (0.3, 0.7).

#### 3.1.3 Coverage

To demonstrate the benefit of multiple imputations, we measured the coverage of an estimated ratio, i.e., how close an estimated ratio is from the true ratio. We simulated data with 50% zeros and estimated ratios of all possible pairs of components to accentuate the effect of multiple imputations. To quantitatively compare MIC to the single imputation methods, we repeated 100 times and computed an empirical standard deviation (SD) for each ratio. We then measured the frequency of estimated ratios falling within the median SD of the oracle values for each method. All methods performed better when the counts of taxa were simulated using ZINB than using ZIDM. Interestingly, C05 had much lower coverages than the other two classical methods, although it performed very similarly to them in terms of mean ratios and mean proportions. When ZINB was used, MIC had a coverage of 69.5%, which is around the expected coverage rate of one SD. When ZIDM was used, it had a little lower than the expected coverage. However, it had a much higher coverage than the other methods, as shown in Table 1. A graphical illustration of the coverage is given in Figure S5.

**Table 1:**
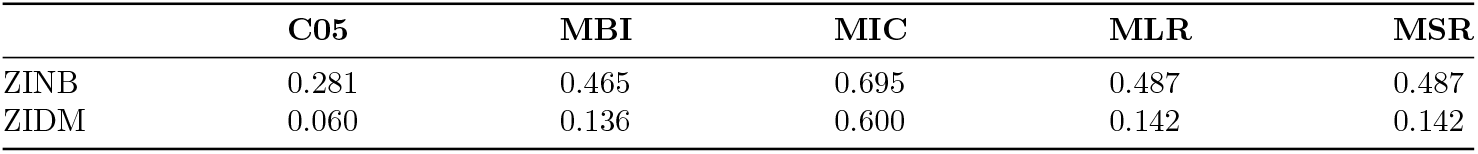
Mean coverage of estimated ratios by the imputation methods. Each value indicates the frequency of estimated ratios within the median SD of the oracle values for each method.

|  | C05 | MBI | MIC | MLR | MSR |
| --- | --- | --- | --- | --- | --- |
| ZINB | 0.281 | 0.465 | 0.695 | 0.487 | 0.487 |
| ZIDM | 0.060 | 0.136 | 0.600 | 0.142 | 0.142 |

#### 3.1.4 Beneficial Effects on Downstream Analysis

We assessed the effects of imputation methods on downstream analysis, specifically differential abundance (DA) analysis, where we identify differentially abundant taxa between groups. Among many statistical methods for DA analysis that account for the compositional effect, we chose LinDA (Zhou *et al*., 2022) because it is fast and performs well in two-group comparisons when the majority of taxa are not differentially abundant, as shown in Sohn *et al*. (2024). We first simulated a taxonomic profile with 50 samples and 100 taxa for each group using ZINB. Model parameters were randomly selected in a setting mimicking real microbiome data, similar to the simulation settings used for mean proportions or ratios. For MIC, we applied LinDA to each of the imputed taxonomic profiles and used Rubin’s rule to pool the results.

Each curve in Figure 6 represents the mean of the receiver operating characteristic (ROC) curve based on 100 repetitions. MBI unexpectedly outperformed the classical methods, although it had lower accuracy (i.e., higher RMSE) in mean ratio and a lower coverage. Also, C05 performed better than MLR and MSR despite its lower coverage, indicating the accuracy in mean ratio and the coverage may not have significant impacts on comparing groups when the counts of taxa are independent of each other, which could also be an explanation of the similar performance of MBI and MIC. However, when a taxonomic profile was simulated using ZIDM, the performance of imputation methods in DA analysis seemed to align with that in accuracy and coverage. MIC outperformed all the methods as it did in terms of accuracy and coverage. As in the accuracy comparison, the performance of MIC was very robust, while other methods were not, demonstrating a beneficial effect of MIC in DA analysis.

**Figure 6:**
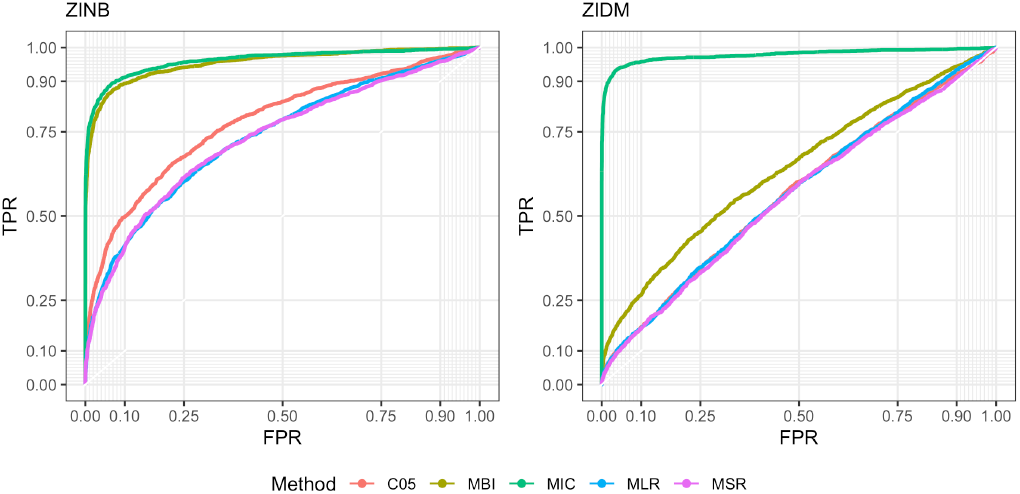
Performance comparison in DA analysis by the receiver operating characteristic (ROC) curve. Each curve is the mean ROC based on 100 repetitions.

### 3.2 Real Data Analysis

We reanalyzed the T2D dataset of Qin *et al*. (2012) available in the R package “curatedMetagenomicData” (Pasolli *et al*., 2017) to determine differentially abundant species between T2D patients and non-diabetic controls. The dataset consists of whole metagenome shotgun sequencing samples of 363 Chinese T2D patients and non-diabetic controls. We removed 15 T2D patients on metformin to avoid any treatment effect on microbial compositions of T2D patients, leaving 193 non-diabetic controls and 155 T2D patients with 155 unique species.

As typical, the taxonomic profile of the T2D dataset was highly sparse, consisting of 51.2% zeros. Using the proposed method MIC, we first generated 200 imputed taxonomic profiles, and then we applied LinDA to each imputed taxonomic profile with the group membership as the primary factor and age, gender, and BMI as covariates. We combined the results using Rubin’s rule. At FDR ≤ 0.2, we identified two differentially abundant species, *Romboutsia ilealis* and *Bacteroides coprocola*, which are all less abundant in T2D patients, as shown in Figure 7. These two species are more abundant in the gut microbiota of healthy individuals, where they regulate blood glucose levels and produce short-chain fatty acids (SCFA), including butyrate, which regulates intestinal physiology and immune homeostasis with an improved insulin response (Chen *et al*., 2017; Rodrigues *et al*., 2021; Baars *et al*., 2024).

**Figure 7:**
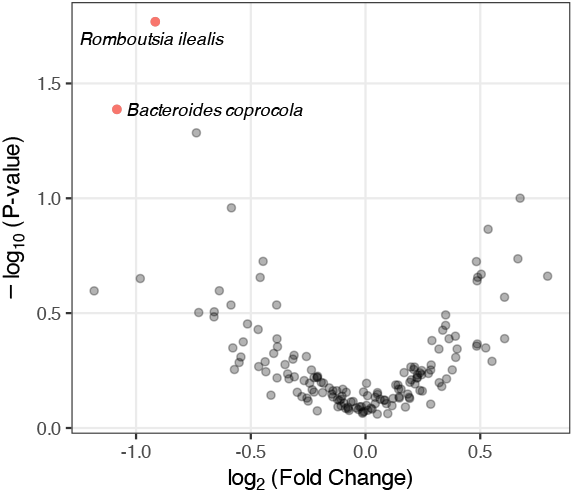
Differentially abundant species between non-diabetic controls and T2D patients. Red dots indicate significantly different species at FDR ≤ 0.2.

## 4 Conclusion

High sparsity is a well-known characteristic of microbiome data, but there is no consensual approach, mainly due to multiple sources of zeros. In this paper, we proposed a semi-parametric multiple imputation method for high-sparse, high-dimensional, compositional microbiome data under the assumption that all zeros are missing values or sampling zeros. We chose this assumption because of the minimal impact of mismodeling structural zeros on composition estimates if the probability of having structural zeros is uniformly distributed around the same mean for all taxa, as empirically demonstrated. However, the proposed approach (MIC) can be adapted to zero-inflated or mixture models just to impute sampling zeros.

The random-selection-and-amalgamation approach implemented in MIC avoids the high sparse and high dimensional issues while capturing some dependence structure in taxa. It also allows for multiple imputations. These properties lend MIC its robust performance over different correlation structures of taxa and a wide range of proportions of zeros, as demonstrated in our extensive simulation studies. Robustness and good coverage are essential attributes for any statistical method since we never know the true structure of real data.

The coverage is controlled by the number of taxa used for the reference in log-ratio. The lower the number of taxa, the higher the coverage. However, a smaller number of taxa leads to a higher probability of having zeros in the reference. Thus, the number of taxa used for the reference should not be too small. In all simulation studies, we used 30% ~ 70% of taxa depending on the proportion of zeros. MIC is also computationally efficient. On MacBook Pro with 2 GHz Quad-Core Intel Core 420 i5 and 16 GB RAM, MIC took about 2 seconds to have 200 imputed taxonomic profiles with 100 samples and 100 taxa, while MBI took about 90 seconds to have a single imputed taxonomic profile.

