## Supplemental File for "A semi-parametric multiple imputation method for high-sparse, high-dimensional, compositional data"

### Appendix

#### A Model Parameters Used in Simulation Studies

We simulated taxonomic profiles using two zero-inflated models with the percent of zeros $\xi_j$ ranging from 30% to 80%. The first model was a zero-inflated negative binomial (ZINB) model:

$$X_{ij} \sim \begin{cases} 0 & \text{with probability } \xi_j \\ NB(\mu_j, \gamma_j) & \text{with probability } 1 - \xi_j \end{cases},$$

where $\mu_j$ and $\gamma_j$ are the mean and size parameter of a negative binomial (NB) model for the $j$-th taxon, whose probability mass function is given by

$$P(X_{ij} = x) = \frac{\Gamma(x + \gamma_j)}{\Gamma(\gamma_j)\Gamma(x+1)} \left(\frac{\gamma_j}{\gamma_j + \mu_j}\right)^{\gamma_j} \left(\frac{\mu_j}{\gamma_j + \mu_j}\right)^{x},$$

thus $\mathbb{E}(X_{ij}) = \mu_j$ and $\mathrm{Var}(X_{ij}) = \mu_j + \mu_j^2/\gamma_j$. The mean counts $\boldsymbol{\mu} = (\mu_1, \ldots, \mu_p)^\top$ were randomly generated mimicking real microbiome datasets: 60% from $\{1, 2, 3\} \times s_i$ (low abundance), 30% from $\{4, \ldots, 10\} \times s_i$ (medium abundance), and 10% from $\{10, \ldots, 100\} \times s_i$ (high abundance), where $s_i$ is the sequencing depth of the $i$-th sample. The size parameter $\gamma_j$ was randomly generated from a uniform distribution over the interval $[0.5, 2]$, or $\gamma_j \sim \mathcal{U}(0.5, 2)$.

The second model used to simulate count data was a zero-inflated Dirichlet multinomial (ZIDM) model:

$$X_{ij} \sim \begin{cases} 0 & \text{with probability } \xi_j \\ DM(n_i, \boldsymbol{\alpha}) & \text{with probability } 1 - \xi_j \end{cases},$$

where $n_i$ is the total count of the $i$-sample and $\boldsymbol{\alpha} = (\alpha_1, \ldots, \alpha_p)^\top$ is a vector of concentration parameters of a Dirichlet multinomial (DM) model, whose probability mass function is given by

$$P(X_{ij} = x) = \frac{\Gamma(\sum_k \alpha_k)\Gamma(n_i + 1)}{\Gamma(n_i + \sum_k \alpha_k)} \prod_{k=1}^{p} \frac{\Gamma(x_k + \alpha_k)}{\Gamma(\alpha_k)\Gamma(x_k + 1)}.$$

Like the mean counts of the NB model, the concentration parameters were randomly generated from 60% from $\{1, 2, 3\} \times s_i$ (low abundance), 30% from $\{4, \ldots, 10\} \times s_i$ (medium abundance), and 10% from $\{10, \ldots, 100\} \times s_i$ (high abundance), where $s_i$ is the sequencing depth of the $i$-th sample. The total count $n_i$ for the $i$-sample was randomly generated from $\{1, \ldots, 20\} \times 1000$.

For differential abundance (DA) analysis, we first randomly selected $5 \sim 30\%$ of taxa and then multiplied them by values generated from a union of two uniform distributions: $\mathcal{U}(0.1, 0.5)$ and $\mathcal{U}(2, 10)$.

### B Results for Accuracy in Mean Proportions

Figures S1 and S2 show results for mean proportions over 200 repetitions when ZINB or ZIDM was used with $n = 100$ samples and $p = 100$ taxa. Like RMSE of the mean ratios, the proposed method was very robust to the proportion of zeros and the correlation structure, and it performed consistently better than the existing methods.

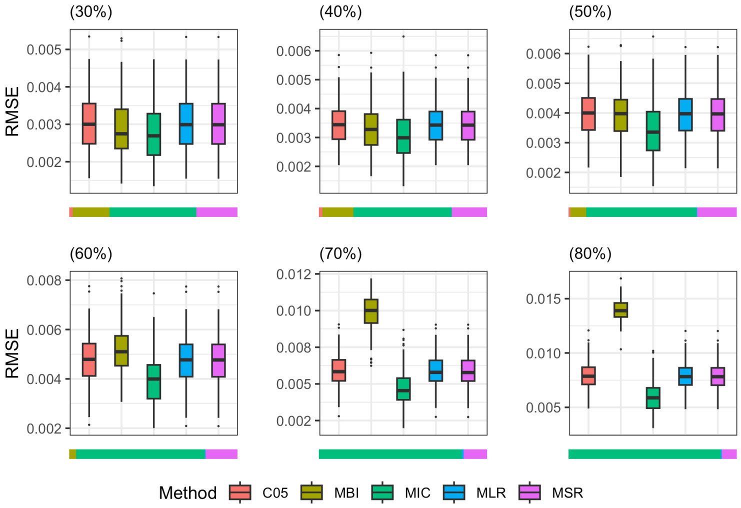


Figure S1: RMSE of Mean Proportions. Zero-inflated negative binomial models were used to simulate data with percent of zeros ranging from 30% to 80%. The horizontal stacked bar plots below the box plots represent the frequency of having the lowest RMSEs over 200 simulations for each method used in the comparison.

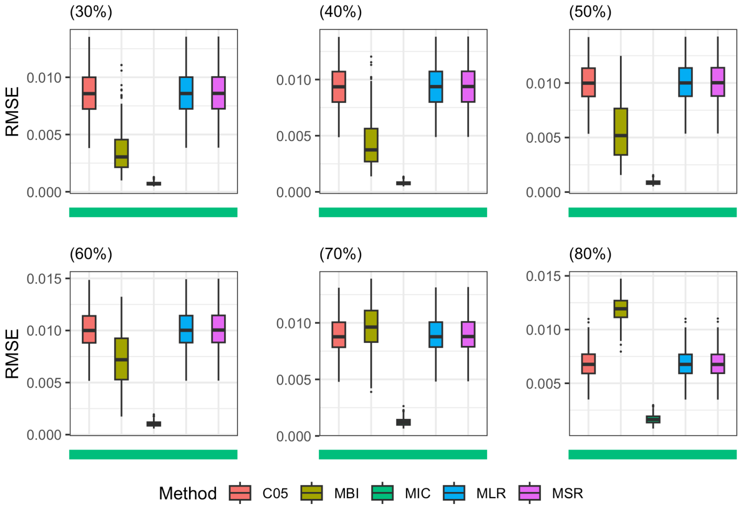


Figure S2: RMSE of Mean Proportions. Zero-inflated Dirichlet multinomial models were used to simulate data with percent of zeros ranging from 30% to 80%. The horizontal stacked bar plots below the box plots represent the frequency of having the lowest RMSEs over 200 simulations for each method used in the comparison.

#### C Results for Effects of Mismodeling Structural Zeros

To investigate the effects of mismodeling structural zeros, we simulated taxonomic profiles using ZINB models with a fixed proportion of zeros for each taxon at 0.5 and a randomly generated proportion of zeros for each taxon from a uniform distribution over the interval (0.3, 0.7). The classical methods were severly affected by mismodeling structural zeros. MBI was less severely affected in terms of mean proportions and not affected in terms of mean ratios. However, its RMSEs in mean ratios seemed to reach the maximum values, thus exhibiting no significant effects of mismodeling structural zeros.

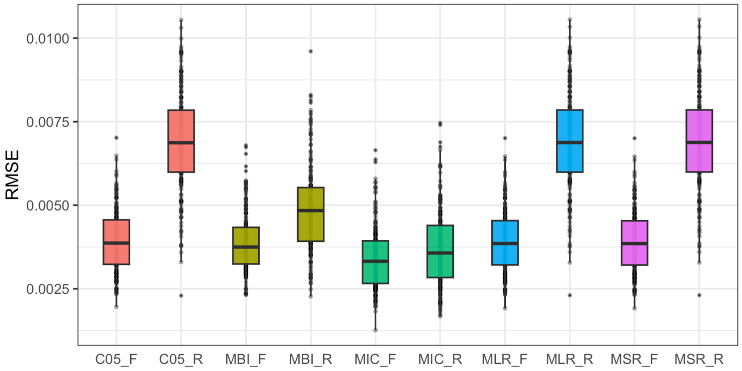


Figure S3: Effects of mismodeling structural zeros on mean proportions. The tailing "F" (e.g., MIC_F) indicates the fixed proportion of zeros at 0.5, and the tailing "R" indicates a randomly generated proportion of zeros from a uniform distribution over the interval (0.3, 0.7).

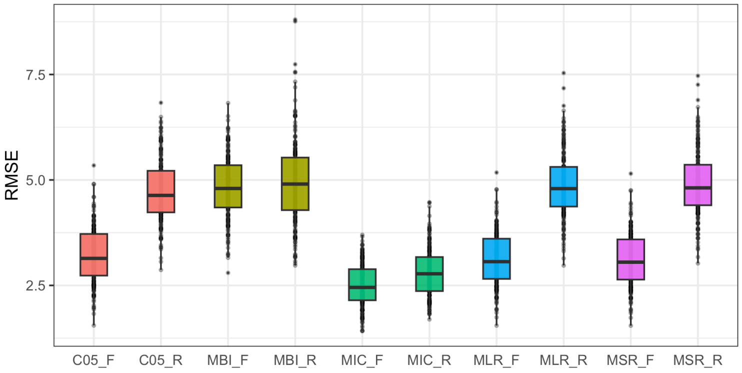


Figure S4: Effects of mismodeling structural zeros on mean ratios. The tailing "F" (e.g., MIC_F) indicates the fixed proportion of zeros at 0.5, and the tailing "R" indicates a randomly generated proportion of zeros from a uniform distribution over the interval (0.3, 0.7).

#### D Coverage

To illustrate a benefit of multiple imputation, we measured the coverage of estimated ratios. Each red point in Figure S5 represents each of ratios estimated using C05, a single imputation method. Each gray vertical line represents a 95% confidence interval of each of ratios estimated using MIC, a multiple imputation

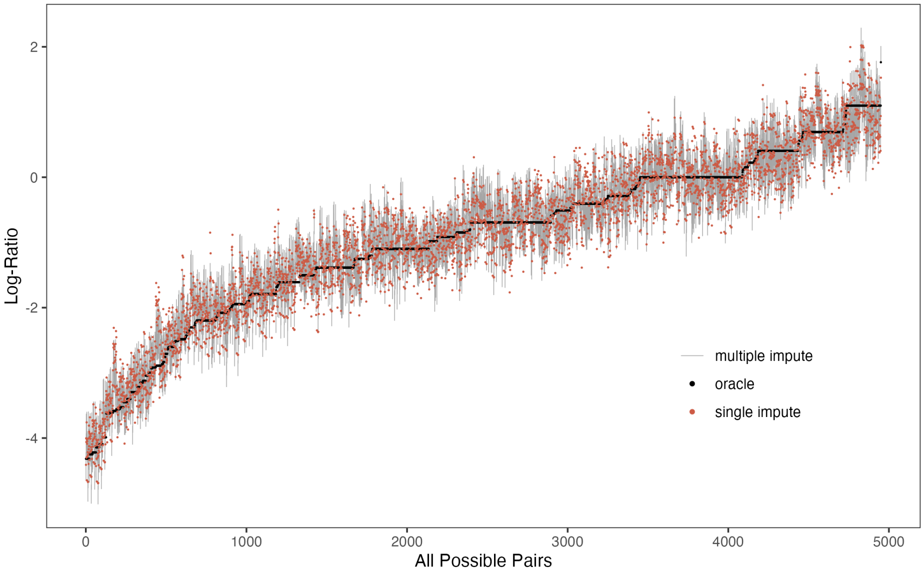


Figure S5: Coverage of estimated ratios. The x-axis indicates ratios constructed with all possible pair of 100 components in order of their log-ratios. The black dots represent the oracle values and the red dots estimated ratios using C05, a single imputation method. The gray vertical lines represent 95% CIs of ratios estimated using MIC, a multiple imputation method.

method. As shown in Figure S5, most red points are far from the true or oracle values although the points scatter around the oracle values, whereas many gray vertical lines include the oracle.
